# Chronic dietary erythritol exposure elevates fasting plasma erythritol levels but does not cause weight gain or modify glucose homeostasis in mice

**DOI:** 10.1101/2021.01.11.426217

**Authors:** Semira R. Ortiz, Martha S. Field

**Affiliations:** Division of Nutritional Sciences, Cornell University, Ithaca, NY 14853, USA

**Keywords:** non-nutritive sweetener, erythritol, obesity, glucose tolerance

## Abstract

**Objective:** Erythritol is both a common non-nutritive sweetener (NNS) and an endogenous product of glucose metabolism. Recent reports indicate that elevated plasma erythritol is a predictive biomarker of cardiometabolic disease onset and complications. Although short-term erythritol consumption has been evaluated, the effect of chronically elevated circulating erythritol on adiposity and glucose metabolism has not. This study investigated the effect of longer-term erythritol consumption on weight gain and glucose tolerance, and the interaction between dietary composition and erythritol supplementation on these parameters.

**Methods:** 8-week-old and 20-week-old C57BL/6J mice were randomized to consume low-fat diet (LFD), high-fat diet (HFD), LFD with 40g/kg erythritol (LFD+ERY), and HFD with 40g/kg erythritol (HFD+ERY) groups. After 8 weeks, plasma erythritol, body weight and composition, food intake, glucose tolerance, and brown adipose tissue (BAT) uncoupling protein 1 (UCP1) expression were measured.

**Results:** Plasma erythritol was elevated 40-fold in mice consuming LFD+ERY or HFD+ERY relative to mice consuming LFD or HFD, respectively. Liver and kidney tissue contained higher levels of erythritol than adipose. Unexpectedly, there was no effect of erythritol supplementation on body weight or glucose tolerance in 8- or 20-week-old mice fed LFD+ERY, or in 8-week-old mice fed HFD+ERY. In 20-week-old mice fed HFD+ERY, there was a significant interaction between erythritol and body weight (p<0.0001) compared to controls, but the main effect of diet was not significant. We also found no effect of chronic erythritol consumption on BAT UCP1 expression.

**Conclusion:** Prolonged erythritol consumption did not significantly impact body weight, composition, or glucose tolerance. This suggests that dietary erythritol does not contribute to the development of cardiometabolic disease.

## Introduction

Reducing obesity rates is imperative due to the severity of obesity comorbidities and the burden chronic disease poses to the healthcare system. Elevated obesity rates are attributed to low physical activity and consumption of a “western” diet characterized by relatively high levels of refined sugar, salt, and saturated fat. Decreasing consumption of sugar is a key point of obesity intervention because added sugars contribute to excess caloric intake, promoting a positive energy balance.

Non-nutritive sweeteners (NNS), zero or low-calorie alternatives to sucrose, are a common alternative to refined sugar because they reduce caloric intake while maintaining the sweet taste of foods. Serum levels of one NNS, erythritol, were recently identified as a predictive biomarker of a range of adverse health outcomes.^1^ The first association between erythritol and metabolic dysfunction found that young adults that gained central adiposity over their first year of college had baseline serum erythritol levels 15-fold higher than those with stable weight over the same time period.^2^ In the larger Atherosclerosis Risk in Communities (ARIC) cohort, serum erythritol was significantly associated with both incident T2DM and cardiovascular disease risk.^3–5^ In these studies, non-targeted serum metabolomics were performed on serum collected at baseline and analyzed for association with cardiometabolic disease onset up to 20 years later. Serum erythritol improved prediction of diabetes onset beyond fasting glucose or other known diabetes risk factors.^3^ Serum erythritol is also associated with disease complications including: diabetic retinopathy, nephropathy, arterial stiffness, vascular complications, and preeclampsia.^6–10^

Erythritol, which naturally occurs in fruits and fermented foods, is a sugar alcohol that is about 70% as sweet as sucrose.^11^ In food production, erythritol is frequently used to sweeten beverages and can be included in nutrition labels as a “natural flavor” at concentrations up to 1.25%. It is also favored as a bulking agent when combined with high intensity sweeteners (e.g. aspartame, sucralose, stevia) because of its crystalline, granulated texture. Erythritol is rapidly absorbed in the small intestine, after which nearly 90% of the erythritol consumed is excreted in urine.^12–14^

In acute doses, studies in humans and rodents have found that erythritol does not impact blood glucose or insulin secretion.^15–18^ The kinetics of erythritol absorption and its effects on postprandial metabolism have been extensively evaluated in short-term studies. There is, however, a paucity of data regarding the impact of prolonged erythritol intake on body composition and glucose metabolism. One relatively longer study using a mouse model of diet-induced obesity found that, over 6 weeks, consumption of a combination of erythritol and aspartame in drinking water increased visceral adipose weight, reduce glucose tolerance, and decrease uncoupling protein 1 (UCP1) expression.^19^ Due to the combined administration of erythritol and aspartame, it is unclear whether erythritol alone produces the same outcome.

Interestingly, Hootman *et. al*. identified that serum erythritol is not only derived from dietary sources. This study demonstrated that erythritol is produced from glucose in humans through the pentose-phosphate pathway (PPP).^2^ Many questions remain regarding endogenous and dietary erythritol. Although dietary erythritol has been shown to be safe during short-term consumption, it has not been determined whether chronically elevated blood erythritol levels are causally related to obesity or cardiometabolic disease. Additionally, it has not been evaluated whether dietary or endogenous erythritol is driving the association between blood erythritol and adverse health outcomes.

The aim of the present study was to determine the effect of prolonged dietary erythritol exposure on body weight, body composition, and glucose metabolism in mice. We fed mice low-fat or high-fat diets supplemented with erythritol for 8 weeks and analyzed the influence of dietary erythritol on body composition, feeding behavior, and glucose tolerance. We also analyzed the impact of diet on endogenous erythritol production and tissue erythritol levels.

## Methods

### Mice

Male C57BL/6J mice were purchased from The Jackson Laboratory at 8 or 20 weeks of age and acclimated to the facility for three days prior to the beginning of dietary treatment. Mice were maintained under specific pathogen-free conditions in accordance with standard of use protocols and animal welfare regulations. All study protocols were approved by the Institutional Animal Care and Use Committee of Cornell University. Mice were housed individually in environmentally controlled conditions (14 hour light/10 hour dark cycle). Mice were randomly divided into one of four dietary treatment groups: 1) low fat (LFD), 2) low fat with 40 g/kg erythritol (LFD + ERY), 3) high fat (HFD), and 4) high fat with 40 g/kg erythritol (HFD + ERY). Each treatment group consisted of N=8 8-week-old mice and N=8 20-week-old mice, for a total of 16 mice per diet treatment. LFD was unmodified AIN-93G formula (Dyets Inc., Bethlehem PA, DYET# 110700GI) with 16% fat-derived calories (FDC). HFD was a modified AIN-93G formula containing 45% FDC (DYET# 104746GI). Erythritol treatment mice were fed either the LFD or HFD formula in which 40 g/kg of cornstarch was replaced by 40 g/kg erythritol (Dyets Inc., Bethlehem PA, DYET#’s 104691GI and 104747GI). 40 g/kg erythritol was chosen to provide the same estimated erythritol consumption as 4% erythritol in drinking water, the dosage observed to induce elevated visceral adiposity by Mitsutomi *et al*.^19^ To mice on LFD and LFD + ERY, food and water were provided *ad libitum* for 8 weeks. All mice in high fat diet groups were placed on HFD for two weeks to induce weight gain. Subsequently, mice in the erythritol group were switched to HFD + ERY for the remaining 6 weeks, while controls were maintained on HFD. All mice were sacrificed after 8 weeks of dietary treatment. An additional N=8 8-week-old mice were fed LFD or HFD for two weeks, then sacrificed, for analysis of tissue erythritol. Intrascapular brown adipose, inguinal white adipose, and epidydimal white adipose tissue were removed, weighed, and frozen in liquid nitrogen. Liver, kidney, quadriceps, and plasma were also harvested and frozen in liquid nitrogen. All tissues were stored at −80°C for use in later applications.

### Food intake, body weight, and body composition measurements

Over the 8-week treatment period, food intake and body weight were measured and recorded twice weekly. Food intake was determined by subtracting the weight of food remaining in the hopper from the weight of food that was supplied. Body composition was measured by NMR after 2 and 8 weeks of treatment using a Bruker Minispec LF65 according to the manufacturer’s protocols. Body composition measurements included free fluid, lean, and fat mass.

### Intraperitoneal glucose tolerance testing

Mice were fasted for 5 hours prior to intraperitoneal glucose tolerance testing (IPGTT). All blood samples were collected from a single nick in the tail vein. Blood glucose was measured via a drop of blood applied to a hand-held glucometer (OneTouch). Prior to glucose injection, fasting blood glucose was measured and a 50 μL blood sample was collected. Blood was collected in an EDTA-coated microvette tube (Sarstedt), centrifuged for 10 minutes at 2,000 x g and 4C, then plasma was transferred to an Eppendorf tube and stored at −80C for later analysis of plasma metabolites. Mice were then injected intraperitoneally with 1.5 mg glucose/g body mass.

Blood glucose was measured at 15, 30, 60, 90, and 120 minutes following glucose injection. The area under the curve was calculated using Prism software.

### Isolation of erythritol from plasma

Polar metabolites were extracted from plasma by combining 20 μL of plasma with 180 μL ice cold extraction fluid consisting of 4:1 methanol and water and 10 μM ^13^C_1_-ribitol (Cambridge Isotopes). Samples were agitated in a Thermoshaker at 4°C for 5 min at 1400 rpm, centrifuged for 5 min at 4°C and 17,000 x g, and 140 μL of the supernatant was transferred to an Eppendorf tube to be dried. The polar phase was evaporated in a vacuum concentrator at −100°C until dry. Erythritol standards ranging from 40 to 0.156 μM erythritol were also extracted as described for plasma erythritol. To the dried sample and standard extracts, 20 μL of 20 mg/mL methoxyamine hydrochloride dissolved in pyridine was added. The extracts were then agitated in a Thermoshaker at 40°C for 90 min at 750 rpm. Next, 20 μL N-methyl-N-trimethylsilyl-fluoroacetamide (MSTFA) was added and extracts were agitated for an additional 30 min at 750 rpm. Derivatized samples and standards were then transferred to a glass vial with inlet for GC-MS measurement.

### Isolation of tissue erythritol

Liver, kidney, quadriceps, and epidydimal adipose tissue were sectioned into 100 mg pieces and place in weighed Eppendorf tube. Ice cold extraction fluid (500 μL/100 mg tissue) consisting of 4:1 methanol and water and 10 μM ^13^C_1_-ribitol was added to each tube of tissue. Next, one 5mm stainless steel tissue homogenizer bead (Qiagen) was added to each tube. Tissues were homogenized in a Qiagen TissueLyser II at max speed for cycles of 30 seconds until tissue was lysed completely. 250 μl/100 mg tissue Milli-Q water was added and vortexed for 10 seconds. 400 μl/100 mg tissue chloroform was then added to the homogenate and vortexed for another 30 seconds. Samples were agitated in a Thermoshaker at 4°C for 15 min at 1400 rpm, centrifuged for 5 min at 4°C and 17,000 x g, and 60 μL of the supernatant was transferred to an Eppendorf tube to be dried. The polar phase was evaporated in a vacuum concentrator at −100°C until dry. Dried extracts were derivatized and transferred to a glass vial with inlet for GC-MS measurement.

### Measurement of erythritol by GC-MS

GC-MS analysis was performed using a Shimadzu GCMS-QP2010 Plus with a Zebron ZB-5MSi capillary column (30 m length, 0.25 mm diameter, 0.25 μm thickness). 1 μL of sample was injected using a Shimadzu AOC-20i auto injector at 270°C in splitless mode. Helium was used as the carrier gas with a flow rate of 0.96 mL/min. The column oven temperature was held at 90°C for 1 min, then increased to 320°C at a rate of 15°C/minute. The oven temperature was held for an additional 8 min at 320°C. The total run time per sample was 24.3 min.

The mass spectrometer was set to electron ionization mode with the ion source held at 240°C and an interface temperature of 300°C. In SIM mode, mass spectra of m/z 217, m/z 307, and m/z 320 were acquired from 8-9 min, followed by m/z 218, m/z 320, and m/z 423 from 10-11 min. The solvent delay was 4.5 min. Erythritol and ^13^C_1_-ribitol peaks were selected from GC-MS chromatograms based on the retention time of their respective standards. Absolute intensities of erythritol (m/z 217) and ^13^C_1_-ribitol (m/z 218) were recorded. The ratio of the absolute intensity of erythritol to that of ribitol for standards and samples was used to determine plasma erythritol concentration. Tissue samples were normalized by dividing the ratio of erythritol to ribitol by tissue mass in grams.

### RNA extraction and quantitative PCR

RNA was isolated from a single lobe of frozen brown adipose tissue using RNA Stat-60 (Tel-Test, Inc.). RNA was reverse transcribed using the High-Capacity cDNA Reverse Transcription Kit (Applied Biosystems) according to the manufacturer’s protocol. cDNA was amplified by real-time PCR with the LightCycler 480 SYBR Green I Master mix (Roche). Qiagen QuantiTect primers were used for detection of UCP1 (QT00097300), 18s (QT02448075), and β-Actin (QT00095242). Relative quantification was performed by creating a standard curve for each target mRNA and for 18s or β-Actin (housekeeping genes) mRNA for each sample. Values are reported as the ratio of the target mRNA to 18s or β-Actin mRNA.

### Western Blot Analysis

Frozen BAT samples were homogenized in lysis buffer containing 15% NaCl, 5 mM EDTA, pH 8, 1% Triton X100, 10 mM Tris-Cl, 5 mM DTT, and 10 μl/mL protease inhibitor cocktail (Sigma Aldrich). Protein concentration of the BAT lysate was determined by Lowry assay.^20^ Equal amounts of protein in SDS-PAGE running buffer were boiled then loaded onto a 10% tris-glycine gel (NuSep) and electrophoresed. Protein was then transferred by electrophoresis to a PVDF membrane (MilliporeSigma). The membrane was blocked in 5% non-fat milk overnight at 4°C, then incubated with primary antibodies (1:1000) against UCP1 and alpha-tubulin (Cell Signaling Technologies) for 1 hour at room temperature. Secondary anti-rabbit antibody (1:10,000, ThermoFisher) was applied to the membrane and incubated for 1 hour at room temperature. Protein was detected using a Protein Simple xxx with Clarity Western ECL Substrate (Bio-Rad). Band intensity was measured using ImageJ (NIH).

### Statistical Analysis

All statistical analyses were conducted using GraphPad Prism 9 (Graphpad Software Inc). Due to missing data, plasma erythritol was analyzed using two-sided unpaired t-tests. The 8-week-old and 20-week-old mice were considered as two separate studies. Bonferroni correction factors of 5 (8-week-old mice) and 3 (20-week-old mice) were applied to account for multiple comparisons. As a result of the Bonferroni corrections, p ≤ 0.01 was considered statistically significant in 8-week-old mice and p≤ 0.016 was considered significant in the 20-week-old group. The effect of diet on tissue erythritol was analyzed using a two-way ANOVA. Differences in body weight over time were assessed with a mixed-effects model (REML) with fixed effects of time, diet, and time x diet. Sidak’s multiple comparisons test was used as post hoc analysis for the mixed-effects model and ANOVA tests to determine differences between groups. The IPGTT area under the curve (AUC) was calculated in GraphPad Prism, then analyzed with a two-sided unpaired t-test. All other data were analyzed using a two-sided unpaired t-test and p≤ 0.05 was considered statistically significant.

## Results

### Erythritol is synthesized endogenously in mice, and fasting erythritol is significantly elevated by erythritol supplementation

To assess the effect of dietary composition on circulating plasma erythritol, we measured plasma erythritol levels in fasted mice exposed to 16% FDC (LFD) or 45% FDC (HFD) diet with or without added erythritol (+ ERY). Plasma erythritol levels were significantly higher in all mice fed 40g/kg erythritol (LFD+ERY and HFD+ERY) compared to their respective control diets (LFD or HFD) at both time points (p<0.001, Table 1). Notably, these 40-fold higher plasma erythritol levels in erythritol-fed mice were present even after a 5-hour daytime fast (Table 1). In addition, endogenous plasma erythritol levels (plasma erythritol in mice consuming LFD or HFD) ranged from 0.3 to 2 μM. We observed that in 8-week-old mice, after two weeks, plasma erythritol appeared higher in mice consuming the HFD compared to LFD (p=0.025, Table 1). This difference did not reach statistical significance after Bonferroni correction (Table 1). As shown in Figure 1, erythritol was present in all tissues measured and was significantly different based on tissue type (p<0.0001) and diet (p<0.05). Liver and kidney contained the highest relative erythritol levels (per gram weight), while only minimal erythritol was present in white adipose. Post-hoc analysis revealed that in the kidney, erythritol was significantly higher in mice fed LFD (Figure 1 B, p<0.05).

**Table 1.**
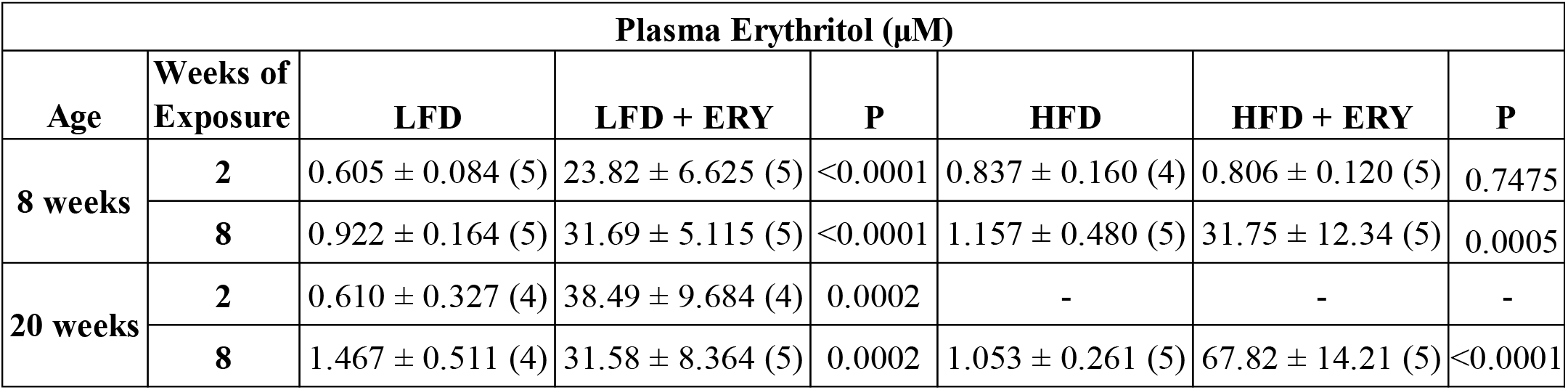
Fasted plasma erythritol levels in 8 and 20-week-old mice fed experimental diets. After 2 or 8 weeks of dietary treatment, plasma was sampled following a 5-hour fast. Erythritol was measured by GC-MS in 8-week-old and 20-week-old mice fed control or erythritol-containing diets. In 8-week-old mice at 2 weeks of exposure, both HFD and HFD+ERY are consuming HFD (without erythritol) to initiate weight gain. Data are presented as mean ± SD (n) and were analyzed by student’s t-test with Bonferroni correction. P-values represent t-test of erythritol vs. respective control diet. P≤0.01 is considered statistically significant to account for a Bonferroni correction factor of 5 in the 8-week group. P≤0.016 was considered significant for the 20-week group accounting for a Bonferroni correction factor of 3. LFD: low fat diet; HFD: high fat diet; LFD + ERY: low fat diet with 40g/kg erythritol; HFD + ERY: high fat diet with 40g/kg erythritol.

| Plasma Erythritol (μM) |  |  |  |  |  |  |  |
| --- | --- | --- | --- | --- | --- | --- | --- |
| Age | Weeks of Exposure | LFD | LFD + ERY | P | HFD | HFD + ERY | P |
| 8 weeks | 2 | 0.605 ± 0.084 (5) | 23.82 ± 6.625 (5) | <0.0001 | 0.837 ± 0.160 (4) | 0.806 ± 0.120 (5) | 0.7475 |
|  | 8 | 0.922 ± 0.164 (5) | 31.69 ± 5.115 (5) | <0.0001 | 1.157 ± 0.480 (5) | 31.75 ± 12.34 (5) | 0.0005 |
| 20 weeks | 2 | 0.610 ± 0.327 (4) | 38.49 ± 9.684 (4) | 0.0002 | - | - | - |
|  | 8 | 1.467 ± 0.511 (4) | 31.58 ± 8.364 (5) | 0.0002 | 1.053 ± 0.261 (5) | 67.82 ± 14.21 (5) | <0.0001 |

**Figure 1.**
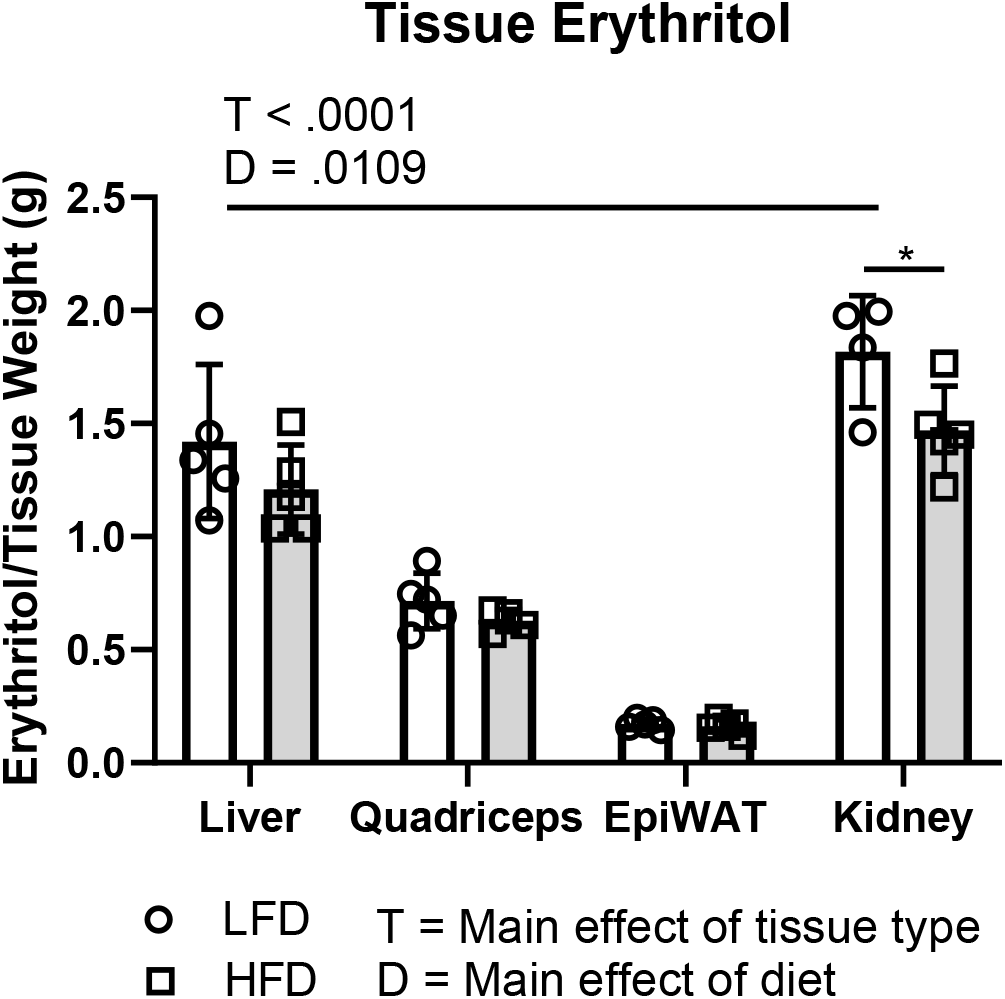
Comparison of endogenously synthesized erythritol in mice fed control diets. 8-week-old mice were exposed to control LFD or HFD (without erythritol) for two weeks. Endogenous tissue erythritol was measured by GC-MS and normalized to tissue wet weight. Data are presented as mean ± SD and were analyzed by two-way ANOVA: *p<0.05. epiWAT: epididymal white adipose tissue; LFD: low-fat diet; HFD: high-fat diet.

### Chronic dietary erythritol consumption does not cause weight gain in mice consuming a low-fat diet

To determine whether dietary erythritol causes weight gain or glucose intolerance outside the context of diet-induced obesity, we fed 8-week-old and 20-week-old mice control (LFD) or erythritol supplemented (LFD + ERY) low-fat diet for 8 weeks. Unexpectedly, there was no difference in body weight (Figures 2A and 3A) or cumulative calorie intake (Figures 2B and 3B) between mice consuming LFD and LFD + ERY. This was true for both young and older mice (Figures 2A, B and 3A, B). Likewise, there was no significant difference in glucose tolerance after 8 weeks dietary erythritol exposure in either young (Figure 2 C, D) or aged mice (Figure 3 C, D), as indicated by the intraperitoneal glucose tolerance test (IPGTT) area under the curve. Similarly, there was no difference between diets in total body fat or adipose tissue weights in either age group (Figures 2 E, F and 3 E, F).

**Figure 2.**
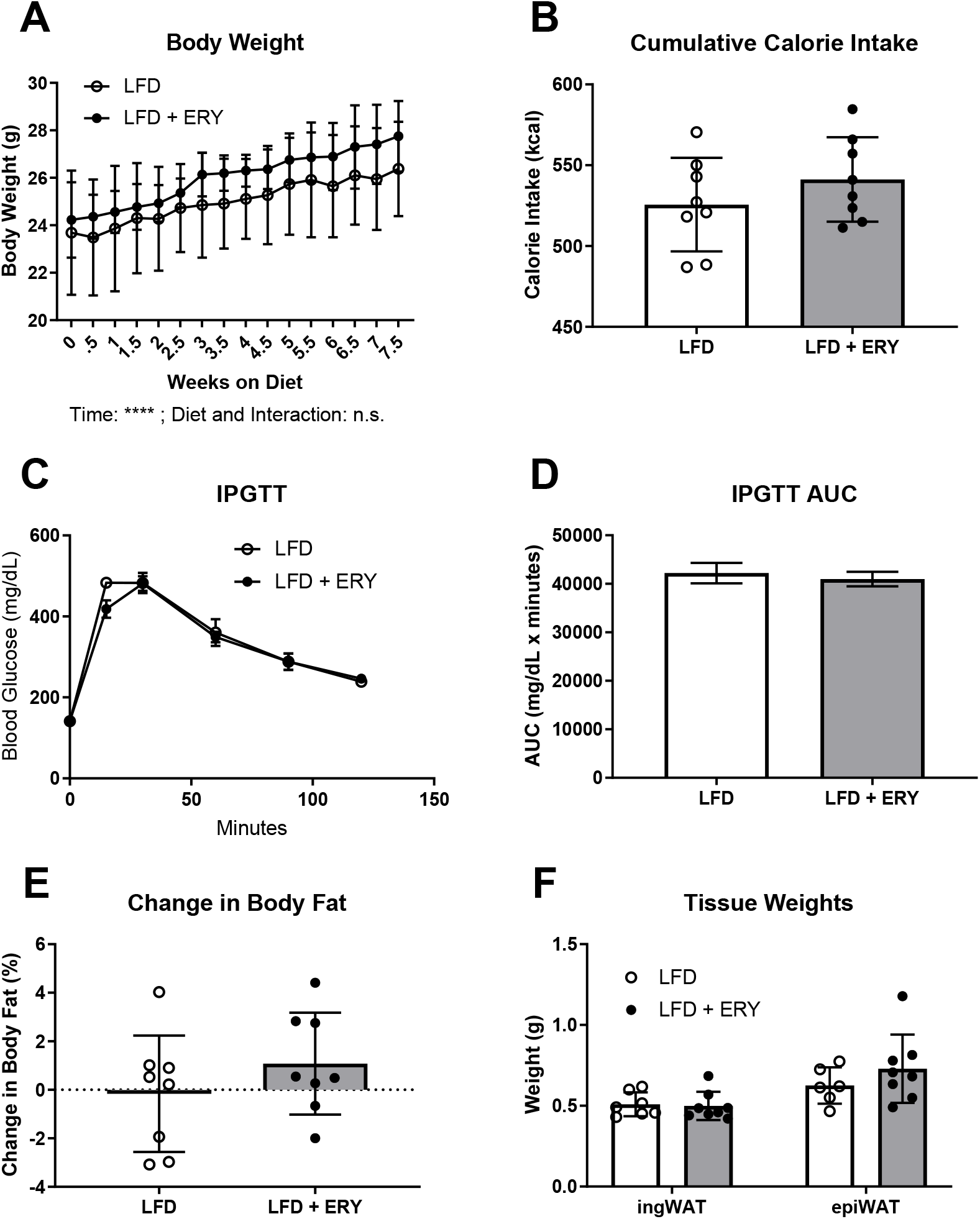
Effect of dietary erythritol on body weight, food intake, and glucose tolerance parameters in 8-week-old mice fed LFD or LFD + ERY. A) Weekly body weight in grams. B) Total caloric intake in kilocalories (kcal). C) Results of intraperitoneal glucose tolerance testing (IPGTT), presented as mean ± SEM, n=5. D) Area under the curve (AUC) of panel C. E) Change in body fat percentage, measured by NMR. F) Adipose depot weights. Data, excluding panel C, are presented as mean ± SD, ****p<0.0001. LFD: low fat diet; LFD + ERY: low fat diet with 40g/kg erythritol; epiWAT: epididymal white adipose tissue; ingWAT: inguinal white adipose tissue.

**Figure 3.**
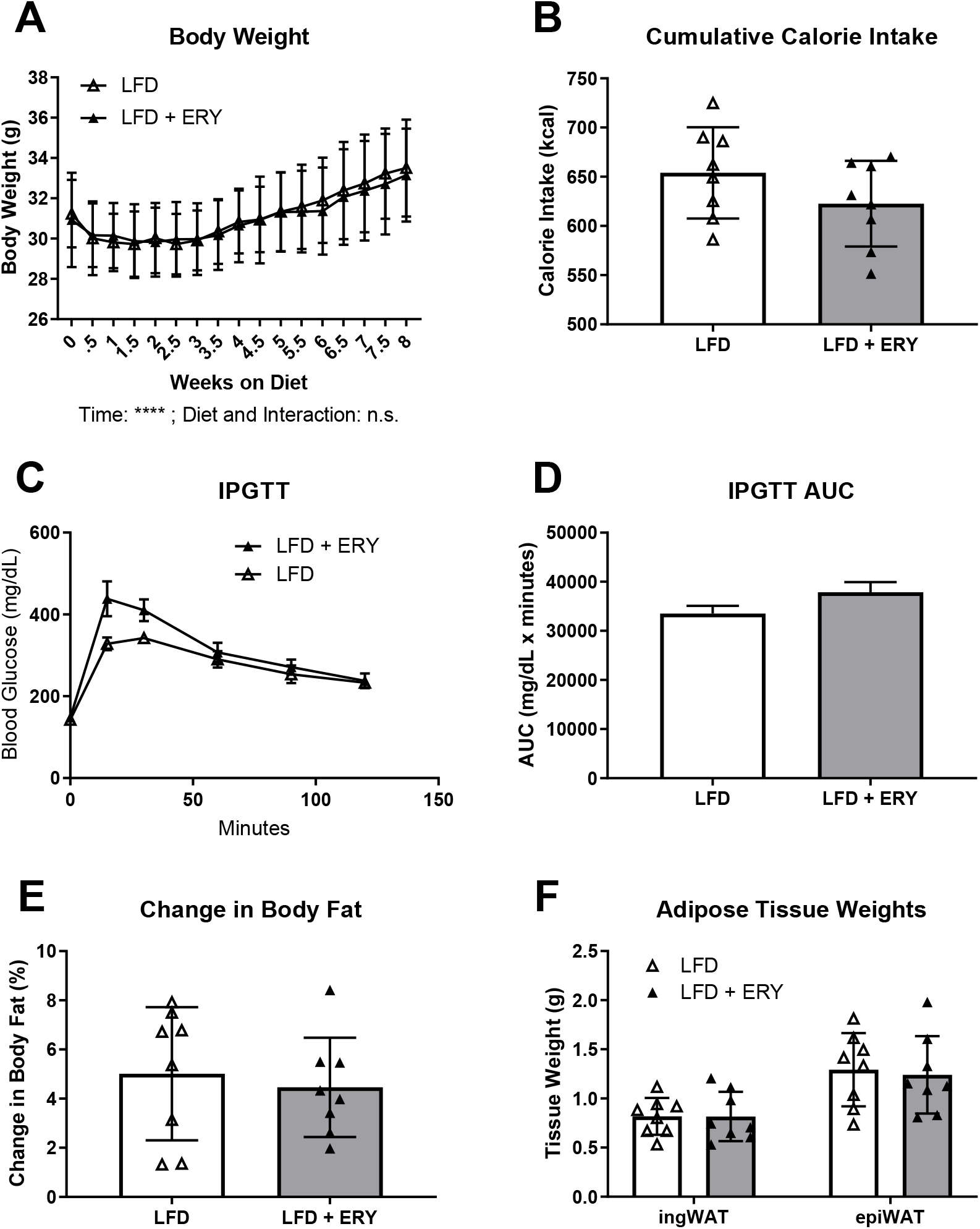
Effect of dietary erythritol on body weight, food intake, and glucose tolerance parameters in 20-week-old mice fed LFD or LFD + ERY. A) Weekly body weight in grams. B) Total caloric intake in kilocalories (kcal). C) Results of intraperitoneal glucose tolerance testing (IPGTT), presented as mean ± SEM, n=5. D) Area under the curve (AUC) of panel C. E) Change in body fat percentage, measured by NMR. F) Adipose depot weights. Data, excluding panel C, are presented as mean ± SD, ****p<0.0001. LFD: low fat diet; LFD + ERY: low fat diet with 40g/kg erythritol; epiWAT: epididymal white adipose tissue; ingWAT: inguinal white adipose tissue.

### Chronic dietary erythritol consumption slightly modifies weight gain in diet-induced obese mice

Based on a previous study in diet-induced obese mice, we hypothesized that high-fat diet and erythritol consumption may interact, exacerbating weight gain and glucose intolerance.^19^ In 8-week-old mice, there was no difference in body weight, calorie consumption, glucose tolerance, or adiposity between groups (Figure 4 A-F). In 20-week-old mice, there was a modest interaction between high-fat diet and erythritol supplementation. We observed higher body weight over time in the HFD + ERY group (Figure 5A). The main effect of time and the interaction between time and diet were significant, but the main effect of diet was not (Figure 5 A, p<0.0001). Upon post-hoc analysis, we found no significant difference in body weight between groups at any individual time point (Figure 5 A). Calorie consumption and glucose tolerance were not significantly different between groups (Figure 5 B-D). There was also no significant difference in total body fat percentage or white adipose tissue weight in HFD + ERY compared to HFD mice (Figure 5 E-F).

**Figure 4.**
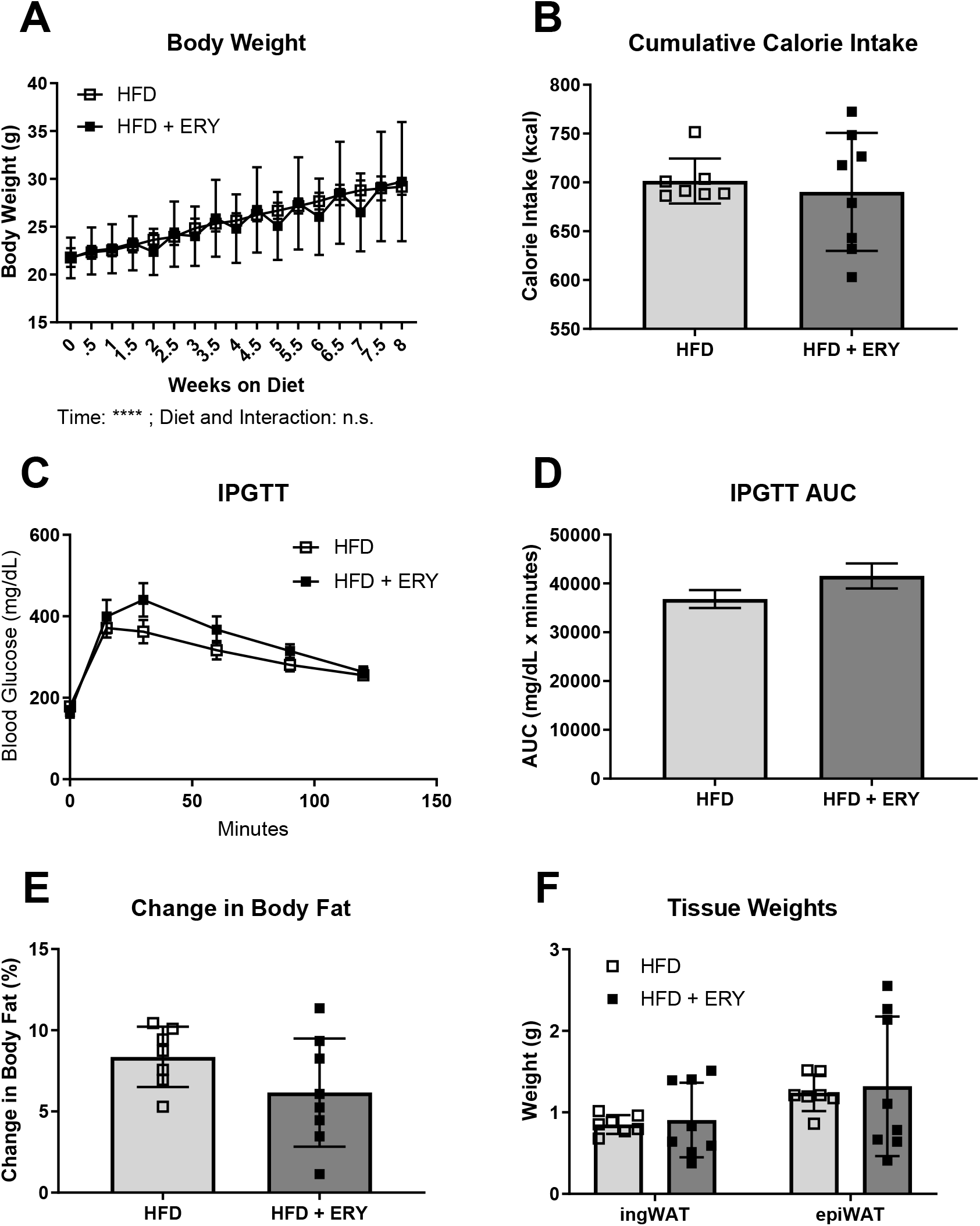
Phenotype data in 8-week-old mice fed HFD or HFD + ERY. A) Weekly body weight in grams. B) Total caloric intake in kilocalories (kcal). C) Results of intraperitoneal glucose tolerance testing (IPGTT), presented as mean ± SEM, n=5. D) Area under the curve (AUC) of panel C. E) Change in body fat percentage, measured by NMR. F) Adipose depot weights. Data, excluding panel C, are presented as mean ± SD. ****p<0.0001. HFD: high fat diet; HFD + ERY: high fat diet with 40g/kg erythritol; epiWAT: epididymal white adipose tissue; ingWAT: inguinal white adipose tissue.

**Figure 5.**
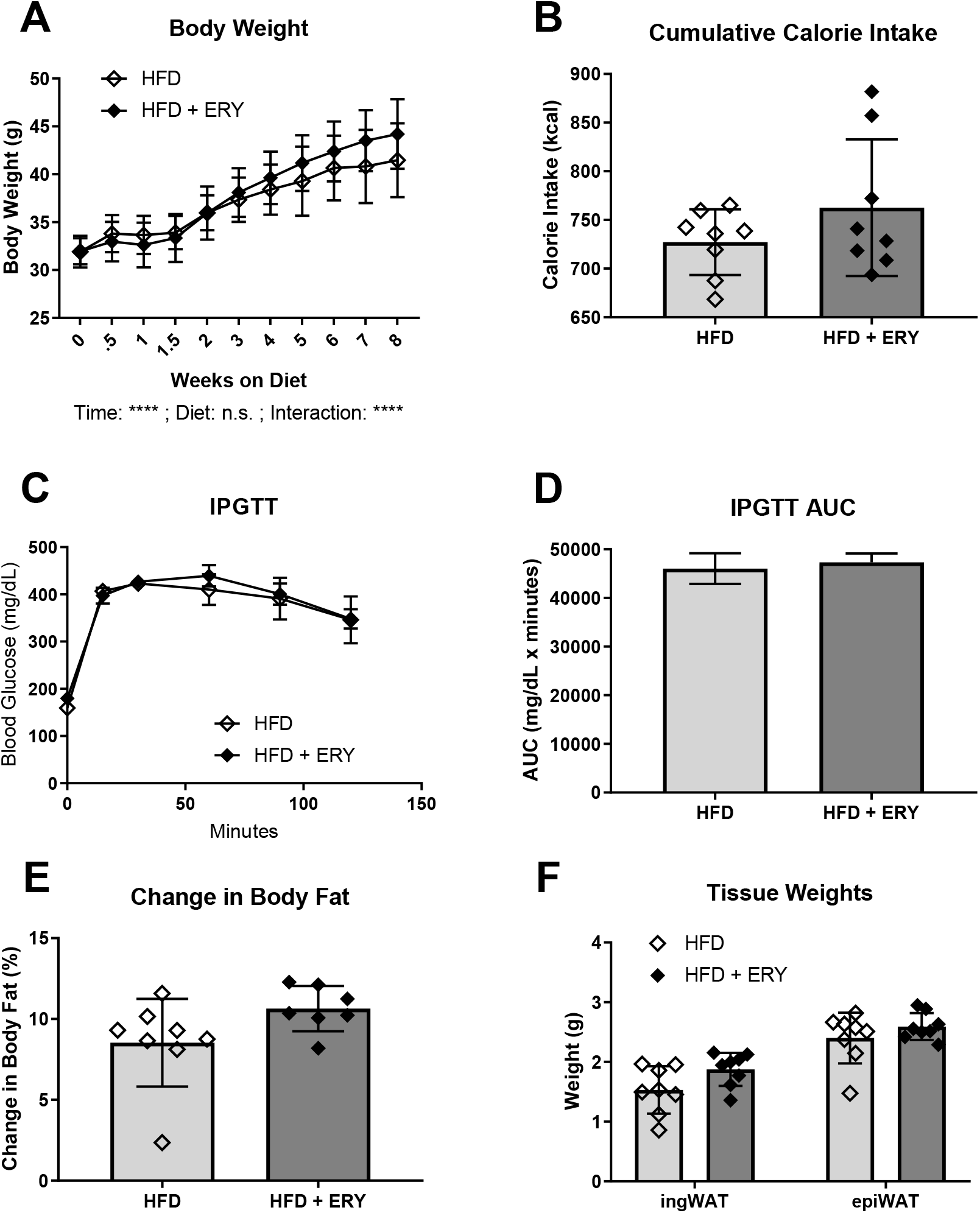
Body composition, food intake, and glucose tolerance phenotype data in 20-week-old mice fed HFD or HFD + ERY. A) Weekly body weight in grams. B) Total caloric intake in kilocalories (kcal). C) Results of intraperitoneal glucose tolerance testing (IPGTT), presented as mean ± SEM, n=5. D) Area under the curve (AUC) of panel C. E) Change in body fat percentage, measured by NMR. F) Adipose depot weights. Data, excluding panel C, are presented as mean ± SD. ****p<0.0001. HFD: high fat diet; HFD + ERY: high fat diet with 40g/kg erythritol; epiWAT: epididymal white adipose tissue; ingWAT: inguinal white adipose tissue.

### Erythritol does not impact UCP1 protein levels in brown adipose tissue

A previous report also demonstrated that in diet-induced-obese mice, non-nutritive sweetener (erythritol and aspartame in combination) consumption significantly reduced BAT UCP1 mRNA expression.^19^ In both age groups of mice fed low-fat diets, there was a significant difference in UCP1 mRNA expression between LFD and LFD + ERY (Figure 6 A and B). The differences in expression, however, were not consistent; in young mice, UCP1 mRNA was significantly decreased, whereas it was significantly higher in 20-week-old animals fed LFD + ERY (Figure 6 A and B, p<0.05). There was also no difference in UCP1 mRNA expression in mice fed HFD or HFD + ERY in either age group (Figure 6 C and D). We additionally assessed protein levels of UCP1 using western blot analysis. There was no difference in BAT UCP1 protein in mice on any dietary intervention (Figure 7 A-C).

**Figure 6.**
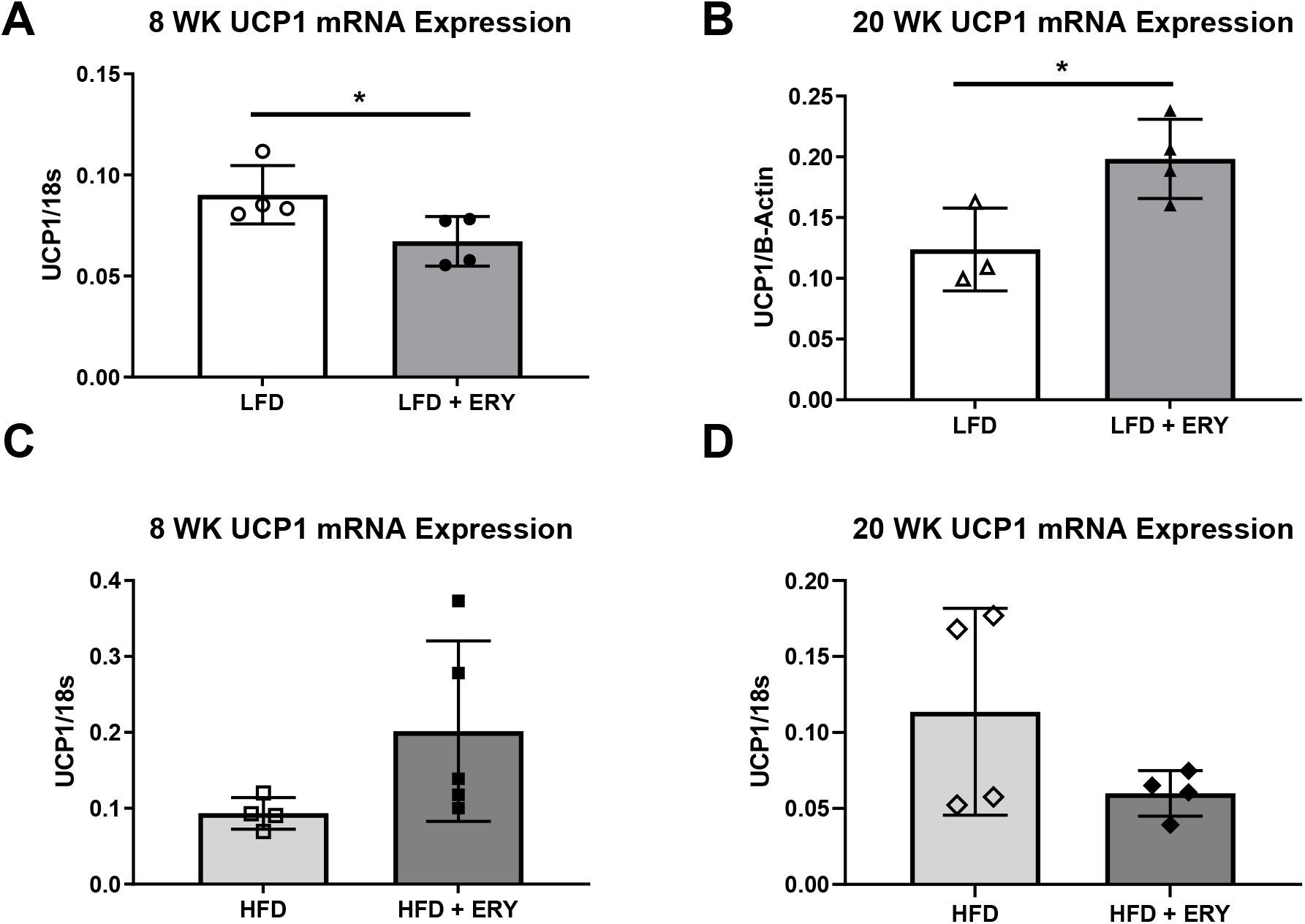
UCP1 mRNA levels in brown adipose tissue of 8 and 20-week-old mice in all groups. Brown adipose UCP1 expression in A) 8-week-old mice and B) 20-week-old mice fed LFD or LFD + ERY or C) 8-week-old mice and D) 20-week-old mice fed HFD or HFD + ERY. UCP1 expression was normalized to 18s or β-actin. Data are expressed as mean ± SD. Differences were analyzed by student’s t-test, n=3-4, *p<0.05. UCP1, uncoupling protein 1; LFD: low fat diet; HFD: high fat diet; LFD + ERY: low fat diet with 40g/kg erythritol; HFD + ERY: high fat diet with 40g/kg erythritol.

**Figure 7.**
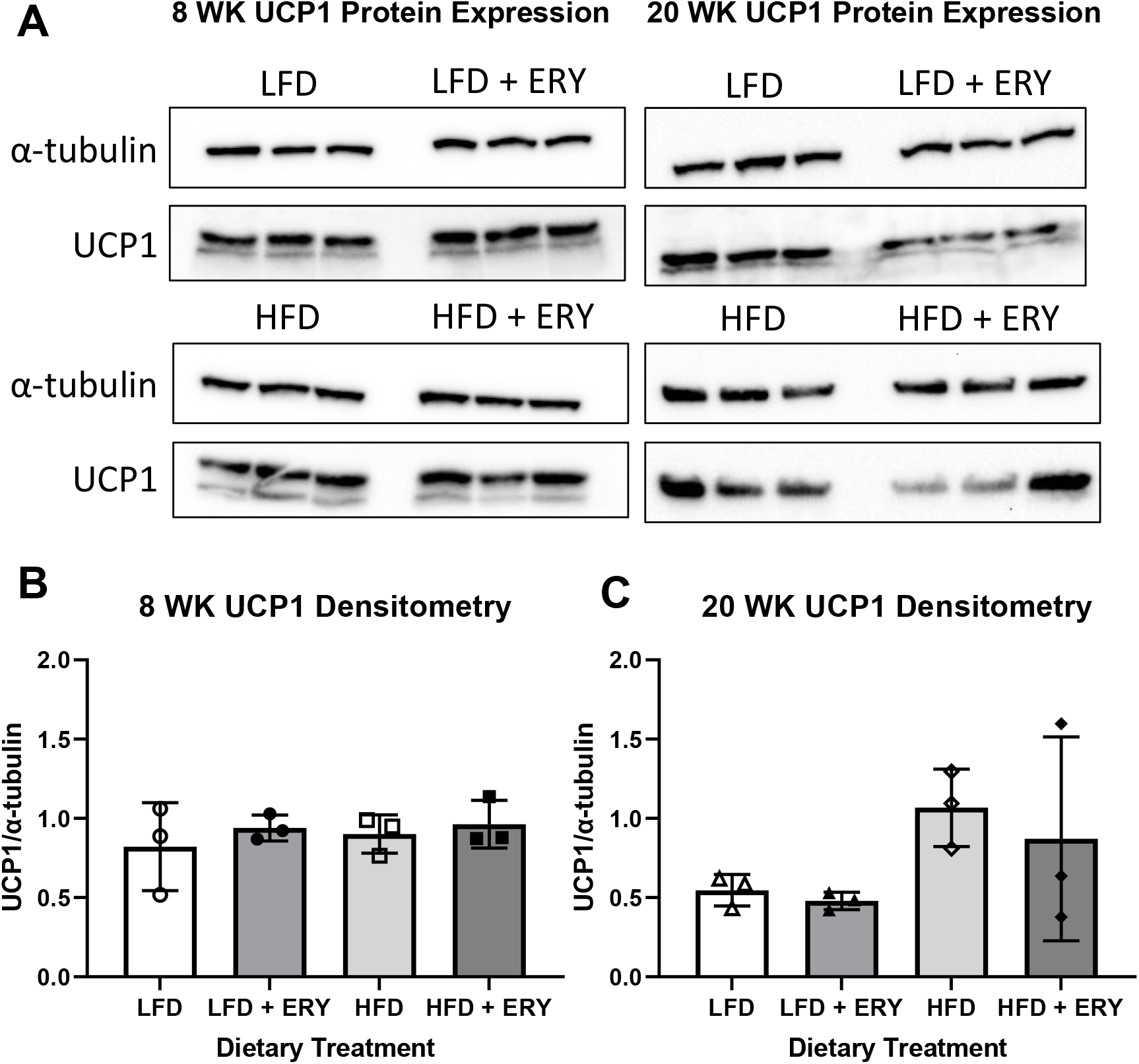
Expression of UCP1 protein in brown adipose tissue of all treatment groups. Protein levels of UCP1 in brown adipose did not differ in 8-week-old (A and B) or 20-week-old (A and C) mice based on exposure to dietary fat or dietary erythritol. Data points represent tissues harvested from individual mice (n=3). UCP1, uncoupling protein 1; LFD: low fat diet; HFD: high fat diet; LFD + ERY: low fat diet with 40g/kg erythritol; HFD + ERY: high fat diet with 40g/kg erythritol.

## Discussion

Elevated plasma erythritol is a predictive biomarker of obesity and diabetes development.^2,3^ Erythritol is both a prevalent sugar substitute and an endogenously synthesized product of glucose metabolism.^1^ Previous studies evaluating the impact of erythritol supplementation on blood glucose and body weight have focused on short-term exposures, most lasting less than two weeks. Here, we characterize the effect of 8 weeks of dietary erythritol consumption on adiposity and glucose homeostasis in mice fed a standard LFD and obesogenic HFD. Given the increasing prevalence of erythritol in the food supply, it is important to understand the effects of chronic erythritol exposure. This is also the first study to describe endogenous erythritol levels in mouse plasma and tissues.

We observed that plasma erythritol was elevated by up to 70-fold over endogenous erythritol levels even after five hours of fasting (Table 1). This is consistent with historic literature assessing the kinetics of erythritol absorption and excretion. Human plasma erythritol peaks within the first several hours of single dose erythritol administration but can remain elevated above baseline up to 8 hours after dosing.^13,21^ Urinary erythritol excretion, the primary fate of dietary erythritol, can continue up to 24 hours after erythritol consumption.^13,21^ Based on this 24-hour timeline, erythritol is considered to be “rapidly” cleared from the body. Here, however, we show that dietary erythritol may increase plasma erythritol levels above baseline/endogenous levels, even under fasted conditions. This suggests that non-nutritive sweetener intake should be carefully considered when assessing the association between plasma erythritol and disease risk.

Surprisingly, in 8-week-old mice, we observed significantly higher plasma erythritol in mice fed HFD compared to LFD (Table 1). While this difference did not reach statistical significance after correction for multiple comparisons (required due to the multiple dietary interventions included in the study), it is worth examining in future studies. Erythritol is synthesized from erythrose-4-phosphate, an intermediate in the non-oxidative PPP.^22^ The non-oxidative PPP consists of a series of interconversions between PPP and glycolytic intermediates, regulated primarily by abundance of glycolytic substrates.^23^ The elevation in plasma erythritol with HFD may be explained by an increase in the availability of glycolytic intermediates, which are funneled into the PPP and then converted to erythritol and secreted to the plasma when energy substrates are in excess. Understanding the relationship between energy balance and increased plasma erythritol levels is outside the scope of the present work.

One limitation of this study is that we were unable to obtain plasma erythritol measurements at the 2-week time point in 20-week-old mice in HFD groups due to global interruptions in research. We expect that these mice would have plasma erythritol within the range of endogenous erythritol we observed in the other three groups.

In tissues, erythritol levels were highest in the liver and kidney, whereas white adipose tissue had relatively low erythritol content (Figure 1). This is the first study to report differential endogenous erythritol levels in metabolic tissues. Interestingly, this pattern of erythritol content parallels relative expression of mouse *Sord* mRNA, one of the enzymes that catalyzes erythritol synthesis *in vitro*.^22^ Mouse liver and kidney have relatively high *Sord* expression, secondary only to the testis, and mouse adipose expresses minimal *Sord* mRNA.^24^ This suggests that SORD contributes to erythritol synthesis *in vivo* in mice. We also found that although plasma erythritol trended higher in mice fed HFD, kidney erythritol levels were significantly, though minimally, lower. It is possible that the differences in plasma erythritol between groups reflect differences in tissue erythritol excretion. As stated above, the dynamics of erythritol excretion under conditions of nutrient excess require further investigation.

In contrast to previous studies, erythritol consumption had no impact on body composition or glucose tolerance in mice consuming LFD.^19^ This indicates that erythritol consumption does not induce metabolic dysfunction in healthy animals. There was also no effect of erythritol on these parameters in young mice fed HFD with erythritol. 20-week-old mice on the same diet, however, trended toward increased body weight, body fat percentage, and adipose weight (Figure 5). This may be attributed to a slight, but not statistically significant increase in food intake. In a previous study, erythritol was used as a calorie control for to D-allulose supplementation in mice fed 40% FDC for 16 weeks.^25^ Compared to 40% FDC mice, the mice fed 50 g/kg erythritol consumed significantly fewer calories and gained less weight.^25^ Trends in body weight appear to follow calorie consumption as opposed to erythritol supplementation. Calorie intake may have been driven higher in HFD + ERY mice due to the hyperpalatable combination of fat and sweetener, which resulted in higher weight and body fat.

As indicated above, Mitsutomi *et al* observed a significant increase in body fat in diet-induced-obese (DIO) mice fed erythritol.^19^ Mice were administered NNS consisting of 99% erythritol and 1% aspartame in drinking water for 4 weeks with 60% fat diet.^19^ Epidydimal white adipose weighed significantly more compared to control DIO mice.^19^ The difference we observed in body fatness of 20-week-old mice fed HFD + ERY compared to HFD was modest compared to the magnitude of response shown by Mitsutomi *et al*. There is evidence that aspartame promotes glucose intolerance and obesity in mice.^26^ The data presented here suggest that the combined effects of erythritol and aspartame, as opposed to erythritol in isolation, in the study by Mitsutomi *et. al*. may have interacted to increase adiposity.

We also assessed BAT *Ucp1* mRNA and protein expression in all groups. Mitsutomi *et. al*. attributed differences in energy utilization between control and NNS-fed mice, in part, to decreased expression of BAT *Ucp1* and its impact on energy expenditure.^19^ Although there were significant differences in *Ucp1* mRNA in mice fed LFD compared to LFD + ERY (Figure 6 A and B), there was no difference in UCP1 protein levels in any group (Figure 7). We, therefore, do not attribute differences in BAT UCP1 to erythritol supplementation.

In summary, prolonged elevation of plasma erythritol through dietary supplementation did not impact weight gain, glucose tolerance, or regulation of energy expenditure via UCP1. Further studies are required to evaluate the relationship between dietary composition, endogenous erythritol, and metabolism.

## Conclusions

In the present study, we did not observe a significant effect of prolonged erythritol consumption on body weight, composition, or glucose tolerance. This suggests that dietary erythritol consumption may not be causally related to cardiometabolic disease development. This is also the first description of endogenously synthesized erythritol in mice. Notably, erythritol was high in liver and kidney, which is consistent with the known expression of SORD (an erythritol-synthesizing enzyme) in these tissues. These findings provide new insight to the relationship between chronically elevated plasma erythritol and the onset of obesity and diabetes.

## Acknowledgements

We would like to thank Stephen Parry and Lynn Johnson with the Cornell Statistical Consulting Unit for their assistance with data analysis and interpretation.

